# Evolution of Delta variant by non-Spike signature co-appearing mutations: trailblazer of COVID-19 disease outcome

**DOI:** 10.1101/2022.04.05.487103

**Authors:** Anindita Banerjee, Anup Mazumder, Jayita Roy, Agniva Majumdar, Ananya Chatterjee, Nidhan K Biswas, Mamta Chawla Sarkar, Arindam Maitra, Shanta Dutta, Saumitra Das

## Abstract

The high transmissibility and infectivity of a SARS-CoV-2 variant is usually ascribed to the Spike mutations, while emerging non-spike mutations might be a serious threat to the current Spike-recombinant vaccines. In addition to mutations in structural Spike glycoprotein, rapid accumulation of mutations across non-structural genes is leading to continuous virus evolution, altering its pathogenicity. We performed whole genome sequencing of SARS-CoV-2 positive samples collected from different clinical groups from eastern India, during the second pandemic wave (April-May, 2021). In addition to the several common spike mutations in Delta variant, two mutually explicit signature constellations of non-spike co-appearing mutations were identified, driving symptomatic and asymptomatic infections. We attempted to correlate these unique signatures of non-Spike co-appearing mutations to COVID-19 disease outcome. Results revealed that the Delta strains harboring a unique constellation of 9 non-spike co-appearing mutations could be the wheeler and dealer of symptomatic infection, even post vaccination. The strains predominantly driving asymptomatic infection possessed 7 non-spike co-appearing mutations, which were mutually exclusive in contrast to the set of mutations causing symptomatic disease. Phylodynamic analysis depicted high probability of emergence of these unique sub-clusters within India, with subsequent spread worldwide. Interestingly, some mutations of this signature were selected in Omicron and IHU variants, which suggest that gradual accumulation of such co-existing mutations may lead to emergence of more “vaccine-evading variants” in future. Hence, unfaltering genome sequencing and tracking of non-Spike mutations might be significant in formulation of any future vaccines against emerging SARS-CoV-2 variants that might evade the current vaccine-induced immunity.

## 1. Introduction

During adaptation to a new host, viruses attempt to make the most of cellular enginery for their efficient replication and protein translation [1]. Viral evolution within an endemic setting frequently encompasses the co-appearance of multiple mutations, either within a single gene or different genes. The effect of single amino acid substitution might be significant to some extent; however, constellation of co-appearing mutations could be necessary to elicit any pathogenicity or difference in disease outcome. Some co-existing mutations are selected as they facilitate viral survival and subsequently predominate in nature [2]. Maximum studies have attributed the high transmissibility and immune-escape properties of the emerging SARS-CoV-2 variants to the Spike structural-glycoprotein mutations [3]. The correlation of mutations in other non-structural proteins (non-Spike mutations) to COVID-19 disease progression is the pressing priority, as this would gauge the effectiveness of the leading Spike protein recombinant vaccines against this deadly virus.

Despite adopting various control measures, COVID-19 second wave had badly hit India with the advent of new variants with increased transmissibility, an uncanny ability of re-infection along with reduced neutralization by convalescent and post-vaccination sera [4-5]. SARS-CoV-2 depicts a strange selectivity with mercurial clinical manifestations among humans [6]. Symptomsrange from mild cold or development of acute respiratory distress syndrome; while few remain asymptomatic. Two possibilities behind these variable symptoms are the virus genotype and the unique host genetic make-up orchestrating their immune responses [7]. In the second wave, several deaths in India occurred due to a “cytokine storm” in adults, while many children contracted the disease with mild to severe symptoms [8]. Multisystem inflammatory syndrome was associated with COVID-19 among children as well as adults [9]. It is indispensable to investigate whether this was completely a host-specific response, or whether SARS-CoV-2 genomic mutations were the key players behind such symptoms.

In addition to host determinants, the co-appearing mutation patterns of the emerging SARS-CoV-2 variants could account for the differences in variable clinical outcomes. Insufficient evidence exists which could associate patients’ infection status (i.e., symptomatic or asymptomatic) to the co-occurring mutations encompassing the viral genome [10]. Maximally, the emerging mutations encompass the Spike glycoprotein, which could modify the binding efficiency to host Angiotensin-Converting Enzyme 2 (ACE2) receptors, alter transmissibility andneutralization efficiency to specific antibodies [11-12]. Nevertheless, mutations across Nucleocapsid and other non-structural proteins might also have a significant impact on viral immune escape strategies, leading to symptomatic or asymptomatic infection [13].

With the aim of tracking the impending SARS-CoV-2 mutations prevailing across eastern India, we attempted to correlate the signature set of non-Spike constellation of mutations and their phylodynamics with the clinical manifestations among both adult and pediatric patients, in the background of their vaccination status. The impact of these mutations might be different due to the variable host immuno-genetic atlas [14-18]. Hence, we have highlighted the predictive biological implications of these co-appearing mutations against some host-proteins involved in inflammatory responses. Correlating the viral non-Spike co-existing mutation patterns with disease outcome will be a prerequisite for designing effective vaccines against emergence of any “vaccine-evading variant” in future.

## 2. Materials and Methods

### 2.1. Clinical sample collection

Nasopharyngeal, oropharyngeal or combined nasopharyngeal/oropharyngeal swabs were collected in Viral Transport Medium (Himedia, MS2760A, India) from patients with clinical features of Influenza like Illness or Severe Acute Respiratory Illness and then sent to the Virus Research and Diagnostic Laboratory (VRDL), Indian Council of Medical Research-National Institute of Cholera and Enteric Diseases (ICMR-NICED), by the state health authorities maintaining cold chain. The Regional VRDL at ICMR-NICED is a Government designated referral laboratory for providing laboratory diagnosis of SARS-CoV-2. At the laboratory, the sample vials were vigorously agitated on a vortex mixture, and the fluid was centrifuged at 1500 rpm for 5 min. This study was based on 244 SARS-CoV-2 positive samples, which were collected during the second pandemic wave in West Bengal (during April-May, 2021) both from unvaccinated pediatric group (below 18 years) (n=60; symptomatic patients=30; asymptomatic ones=30) and adult patients (above 18 years) comprising of 1st dose vaccinated ones (n=50; symptomatic=35, asymptomatic=15), 2nd dose vaccinated (n=42; symptomatic=31, asymptomatic=11) as well as unvaccinated individuals (n=92; symptomatic=45, asymptomatic=47).

### 2.2. Viral RNA extraction and determination of SARS-CoV-2 positivity

Isolation of viral RNAs from the samples was performed using the QIAamp Viral RNA Mini Kit (Qiagen, 52906, Germany), according to the manufacturer’s instructions. SARS-CoV-2 positive samples were detected by quantitative real-time PCR assays. WHO recommended ICMR-NIV Multiplex Single Tube SARS CoV-2 Assay kit (version 3.1) was used. Three SARS-CoV-2 genes, viz., E gene, ORFlab and RdRP, along with one housekeeping gene B-Actin was targeted for amplification. ABI 7500 system was set to capture fluorescence in the range as follows: 520 for E gene (FAM), 550 for ORF (VIC), 580 for RdRP (NED) and 617 for B-Actin (JUN). The cycle threshold was considered as 35 and samples showing signals beyond it were considered negative. The result was interpreted as SARS-CoV-2 positive when the screening gene (E) and any one or both the confirmatory genes (ORF1ab and RdRP) were detected.

### 2.3. Whole genome sequencing

Whole genome sequencing of all the samples included in this study were performed using any one of the following approaches, i.e., paired end sequencing of total RNA libraries post ribosomal RNA depletion by Truseq stranded total RNA library preparation kit (Illumina); genome amplification by standard Arctic primers mainly using COVID-Seq kits (Illumina) or Qiaseq (Qiagen) using Novaseq or Miseq platforms (Illumina). The sequences were submitted to GISAID (https://www.gisaid.org/), bearing accession numbers from EPI_ISL_7210853 to EPI_ISL_7211102 for a total 235 samples.

### 2.4. Phylogenetic analyses

Genome sequences of reference SARS-CoV-2 strains (collected from GISAID and NCBI) belonging to different clades/variants circulating worldwide as well as representative strains from the study were aligned by multiple alignment program MAFFT version 7. Phylogenetic analysis was done using Molecular Evolutionary Genetics Analysis (MEGA) version XI by maximum-likelihood method using the best fit statistical model general time reversal (GTR) with 1000 bootstrap replicates. We also performed the phylogenetic analysis of the representative study strains with the Ultrafast Sample Placement of Existing Trees (UShER) that has been integrated in the UCSC SARS-CoV-2 Genome Browser (https://genome.ucsc.edu/cgi-bin/hgPhyloPlace). UShER is a program that rapidly places new samples onto an existing phylogeny using maximum parsimony.

### 2.5. Comparison of mutated and unmutated SARS-CoV-2 protein-host protein docking interaction

A study on interactome has identified host proteins which physically associate with unmutated SARS-CoV-2 proteins [19]. We have filtered some host proteins that play a crucial role in inflammatory responses. The 3D structure of SARS-CoV-2 proteins (both original and mutated) was generated through RaptorX (http://raptorx.uchicago.edu/) software [20]. 3D structure of the host proteins was obtained from PDB (https://www.rcsb.org/). To predict protein-protein interaction we have used the geometry-based molecular docking algorithm: PatchDock (https://bioinfo3d.cs.tau.ac.il/PatchDock/) [21]. We found different geometric shape complementarity scores for original and mutant types against each viral and host protein interaction. Further, we have refined the PatchDock output using the FireDock (https://bioinfo3d.cs.tau.ac.il/FireDock/) web server where the global energy value was depicted [22]. The more negative (lower) is the global energy value, higher is the binding affinity (less is the binding energy) of the two proteins.

### 2.6. Statistical significance

The statistical significance of the association of the co-appearing mutations with respective clinical groups was evaluated using Fisher’s exact test and the set of mutations that were foundto be significant (p-value ≤ 0.05) has been shown in supplementary figure 1.

## 3. Results

### 3.1. High frequency of SARS-CoV-2 Delta VoC during the second pandemic wave across eastern India, irrespective of the vaccination status

Analyses of the whole genome constellation of 239 SARS-CoV-2 strains (5 out of 244 strains generated low coverage sequences, hence were not considered) collected during April-May, 2021 from West Bengal, eastern India (the time period when India was hard hit by the 2nd wave of COVID-19 pandemic), revealed the preponderance of B.1.617.2 lineage, i.e., the Delta variantof concern (VoC) (n=222/239, 92.8%). Both the pediatric as well as adult patients were predominantly infected by Delta VoC, irrespective of the vaccination status. Some patients showed symptoms of COVID-19 disease and the rest of them were mostly asymptomatic.

In the unvaccinated pediatric group, the overall prevalence of Delta was 91.6%. Susceptibility of Delta infection increased slightly with age (85% among 0-5 years, 90% in 6-10 years, while 100% in 11-15 years). Nevertheless, strains of other lineages like B.1 (20% among ≤10 years), B.1.459 (10% in 6-10 years) and B.1.617.1 (Kappa Variant of Interest; 20% amongst 0-5 yrs children) were also found in traces (Figure 1a). In the adult population, 92.5% infection was Delta mediated. The vulnerability to Delta was slightly less among the vaccinated group (89.6%)in contrast to the unvaccinated ones (95.5%). Other lineages were found in traces, and revealed almost equal frequencies among the vaccinated and unvaccinated groups, i.e., B.1 (2.2% in vaccinated, while 2.1% in unvaccinated group), B.1.459 (1.5% in vaccinated, while 2.1% in unvaccinated group), B.1.617.1 (Kappa VoI; 5% in vaccinated, while 4.5% in unvaccinated group), while B.1.153 lineage was only seen in the vaccinated adults (1.5%) (Figure 1b).

**Figure 1.**
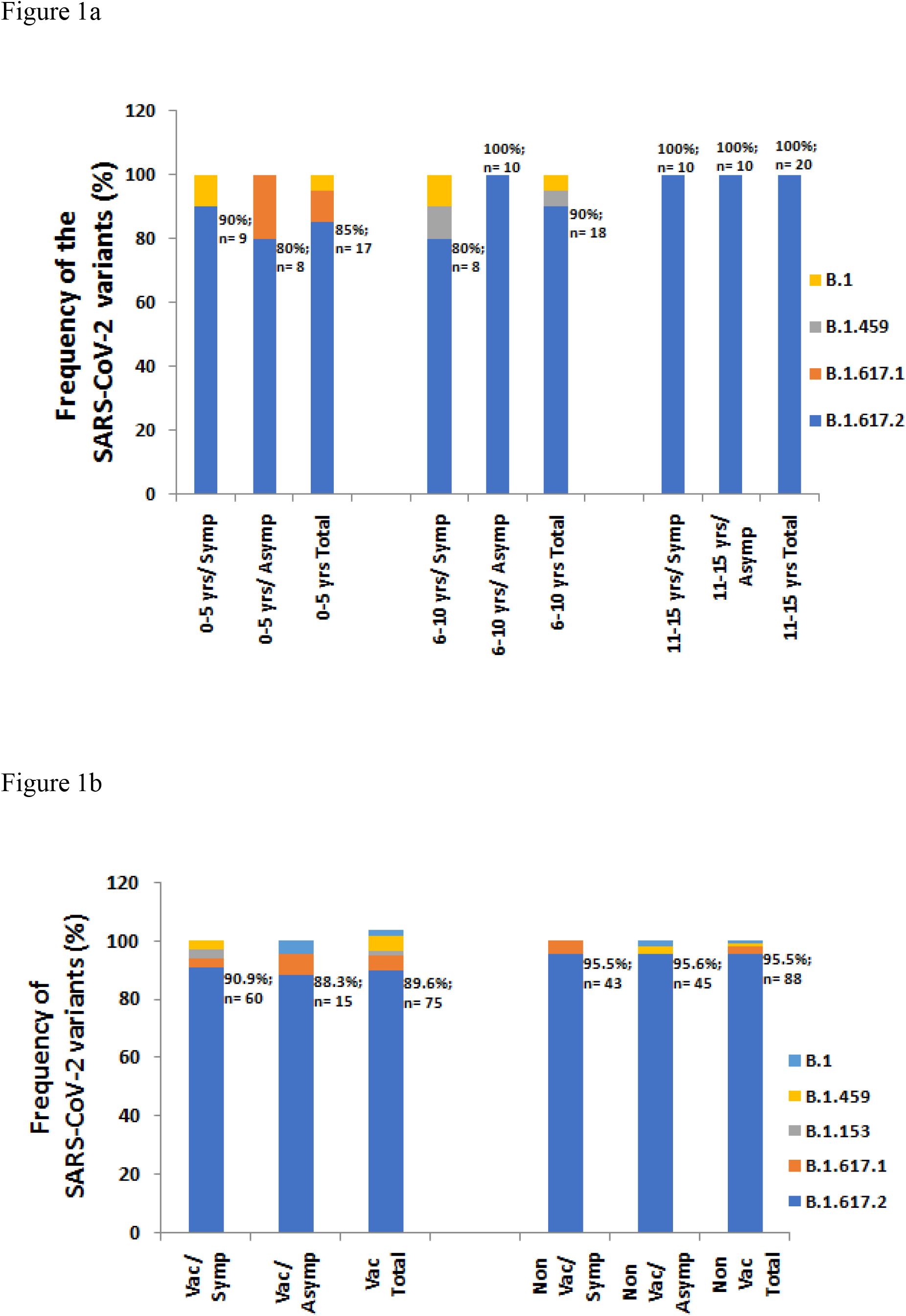
High frequency of the SARS-CoV-2 Delta VoC during the second pandemic wave across eastern India among the (a) pediatric age-group (below 18 years), and (b) adult patients (above 18 years). The unvaccinated pediatric patients have been divided into age-groups like 0-5 years, 6-10 years and 11-18 years, each category sub-divided into symptomatic and asymptomatic groups.The adult patients have been categorized into vaccinated (1st and 2nd dose) as well as unvaccinated groups, where both the categories have been re-divided into symptomatic and asymptomatic groups.

Hence, our initial analysis revealed that Delta VoC was the key player driving the detrimental 2nd COVID-19 wave across eastern India among all age groups, even in the vaccinated individuals and causing symptomatic or asymptomatic disease manifestation.

### 3.2. Phylogenetic analyses revealed significance of additional ORF1a co-appearing mutations in 21J and 21I Delta sub-clade, in eliciting variable disease outcome

Phylogenetic analysis through MEGA software was done for 20 representative Delta strains included in the study (10 from symptomatic clinical group, while another 10 strains from asymptomatic patients) along with multiple SARS-CoV-2 reference genomes (downloaded from GISAID and NCBI GenBank). The representative strains included in the dendrogram were selected based on nucleotide sequence homology among themselves. 10 random representative strains from similar group of strains (with greater than 98% nucleotide sequence homology) were selected. The dendrogram revealed that the 10 representative strains from symptomatic patients sub-clustered with 21J Delta sub-clade strains (sub-cluster-1), while in addition to 21J clade-specific mutations, our strains harbored 2 additional ORF1a co-existing mutations (P2046L, P2287S). The 10 asymptomatic strains formed another distinct sub-cluster 2. Interestingly, this sub-cluster 2 was unique in context to 3 ORF1a co-appearing mutations common to 21I Delta sub-clade (i.e., P1640L, A3209V, and V3718A) (Figure 2a). The representative asymptomatic strains were more homogeneous in context to nucleotide sequences as compared to the strains leading to symptoms. Sporadic synonymous nucleotide substitutions could have led to theheterogeneity among the symptomatic sub-cluster.

**Figure 2.**
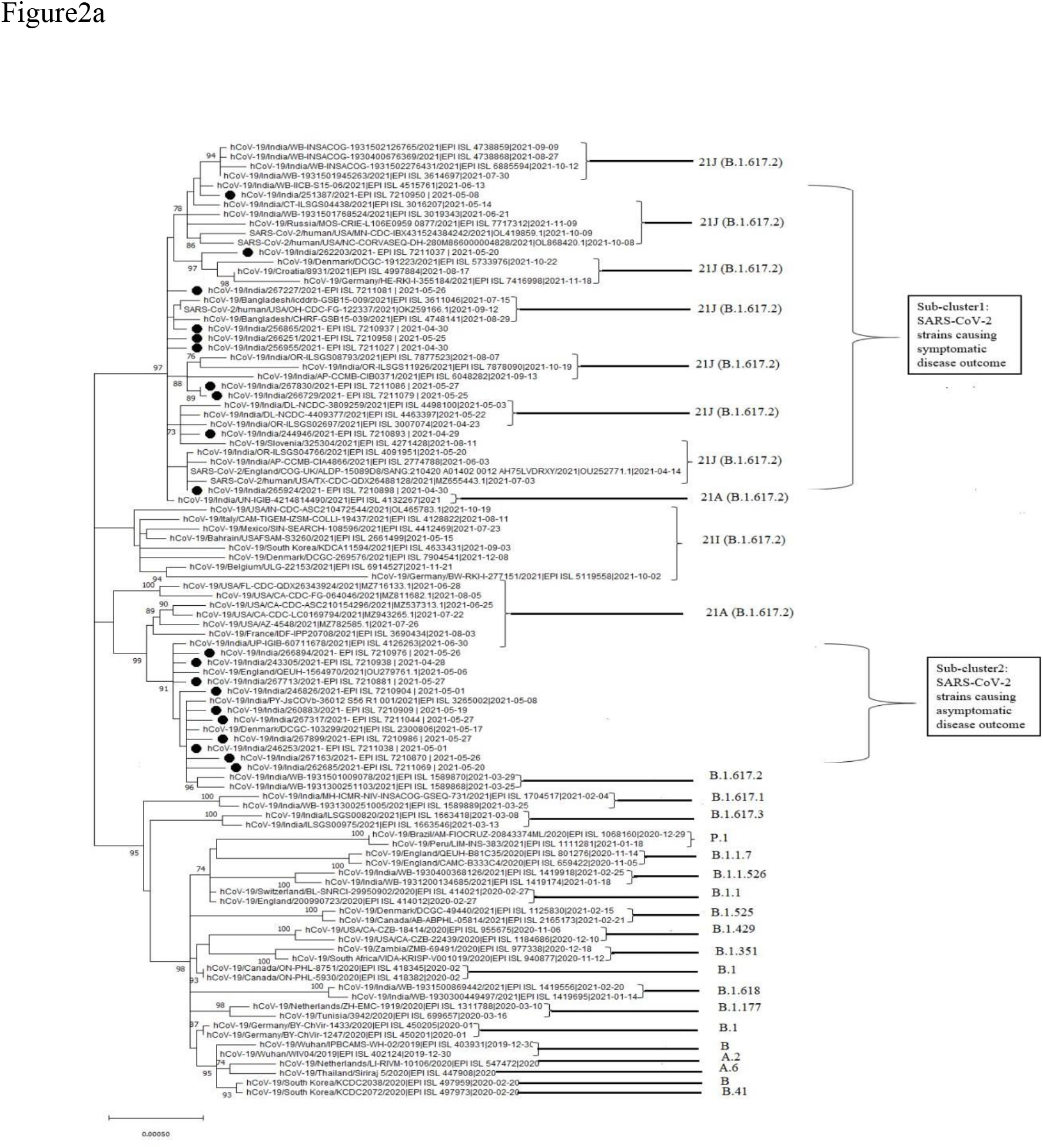
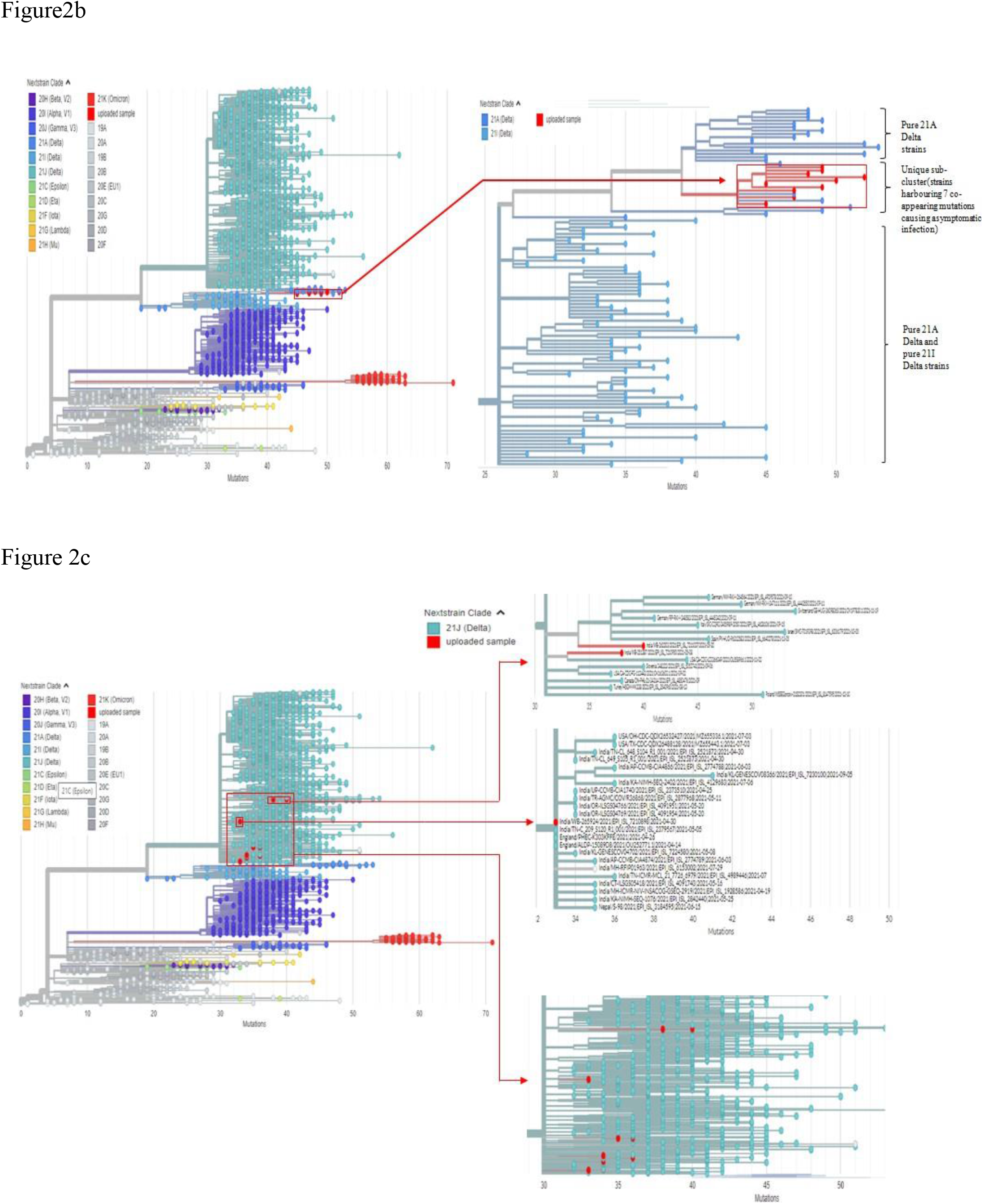
Molecular phylogenetic analysis of the representative Delta strains by (a) MEGA XI using maximum-likelihood method, based on whole genome sequences of 20 representative strains and reference strains belonging to various clades/variants. The scale bar represents 0.00020 nucleotide substitution per site. The best-fit model used for constructing the phylogenetic dendrogram was the general time-reversal model (GTR), (b) UShER, depicting the position of the asymptomatic Delta strains (labelled as uploaded sample in red). Each color is representing a different clade/lineage. Enlarged view of that branch of phylogenetic tree containing the 10 asymptomatic infection causing strain, (c) UShER, depicting the position of the symptomatic Delta strains (labelled as uploaded sample in red). Each color is representing a different clade/lineage. Enlarged view of that branch of the phylogenetic tree containing the 10 symptomatic infection causing strain.

The clustering of all the 20 representative Delta strains was re-analyzed against an already existing dendrogram consisting of 7,217,299 SARS-CoV-2 genomes from GISAID, GenBank, COG-UK and CNCB (submitted till 24^th^ January, 2022) in UShER (https://genome.ucsc.edu/cgi-bin/hgPhyloPlace). Consistent with the results through MEGA XI, all the 10 representative asymptomatic strains formed an extremely unique sub-cluster in the tree generated through UshER. This distinct sub-cluster was neither pure 21A Delta lineage, nor it had all co-existing mutations of 21I Delta sub-clade, while it harbored a signature set of 7 co-appearing mutations (viz., P309L in NSP2, P1640L and H2092Y in NSP3, A3209V in NSP4, V3718A in NSP6, H2285Y in NSP15 and R385K in Nucleocapsid), identified through our analyses (Figure 2b). Only 5 such strains have been submitted in GISAID, all collected post to our sample collection dates [4 from India and 1 from Europe (Denmark)]. While the other 10 representative symptomatic strains, which possessed a signature set of 9 co-appearing mutations (viz., A1306S, P2046L, P2287S in NSP3; V2930L, T3255I in NSP4; T3646A in NSP6; A1918V in NSP14; T40I in ORF7b and G215C in N) clustered with such 21J-like strains from Europe (like Sweden, Norway, England), Asia (India, Bangladesh, Japan) and USA. Collection dates of all such strains from other countries were after our study strains (Figure 2c).

Analyses of the temporal and spatial distribution of the strains with distinct signature sets of co-appearing mutations, leading to differential infection status during 2nd wave, revealed high probability of their emergence in a country like India which harbors a vast population with diverse immuno-genetic atlas.

### 3.3. Emergence of two mutually exclusive signature co-mutation patterns within non-Spike proteins those were associated with symptomatic vs asymptomatic disease outcome

The identified 9 co-appearing mutations (viz., A1306S, P2046L, P2287S in NSP3; V2930L, T3255I in NSP4; T3646A in NSP6; A1918V in NSP14; T40I in ORF7b and G215C in N) were seen to be prevalent (1.96 times higher) among the symptomatic group (26.7%, 27/101); while comparatively less frequent in asymptomatic ones (13.6%, 9/66) (Figure 3a). For the pediatric patients, no difference in frequency was observed among symptomatic and asymptomatic groups (22.2% and 21.4%) (Figure 3b). Except P2046L and P2287S in NSP3, all the 9 co-existing mutations, driving symptomatic infections even after vaccination, were present in the 21J sub-clade of Delta. P2287S is one of the mutations present in Lambda Variant of Interest (VoI), while P2046L was not present in any VoCs or VoIs to date. Surprisingly, strains bearing these array of 9 co-appearing mutations depicted negligible lowering of frequency (1.8%) among the symptomatic-vaccinated individuals (27.6%) in comparison to symptomatic-unvaccinated ones (29.4%).

**Figure 3.**
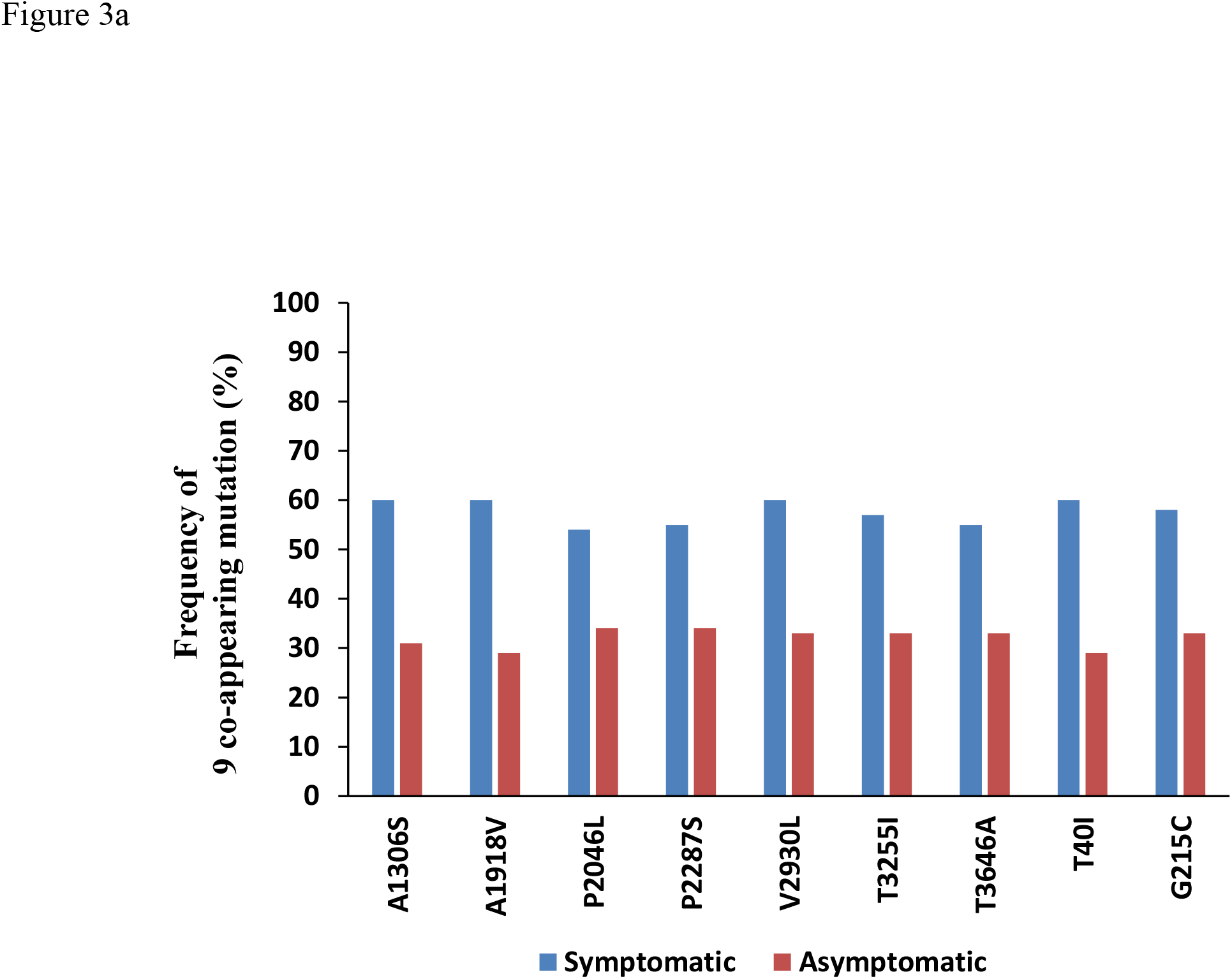
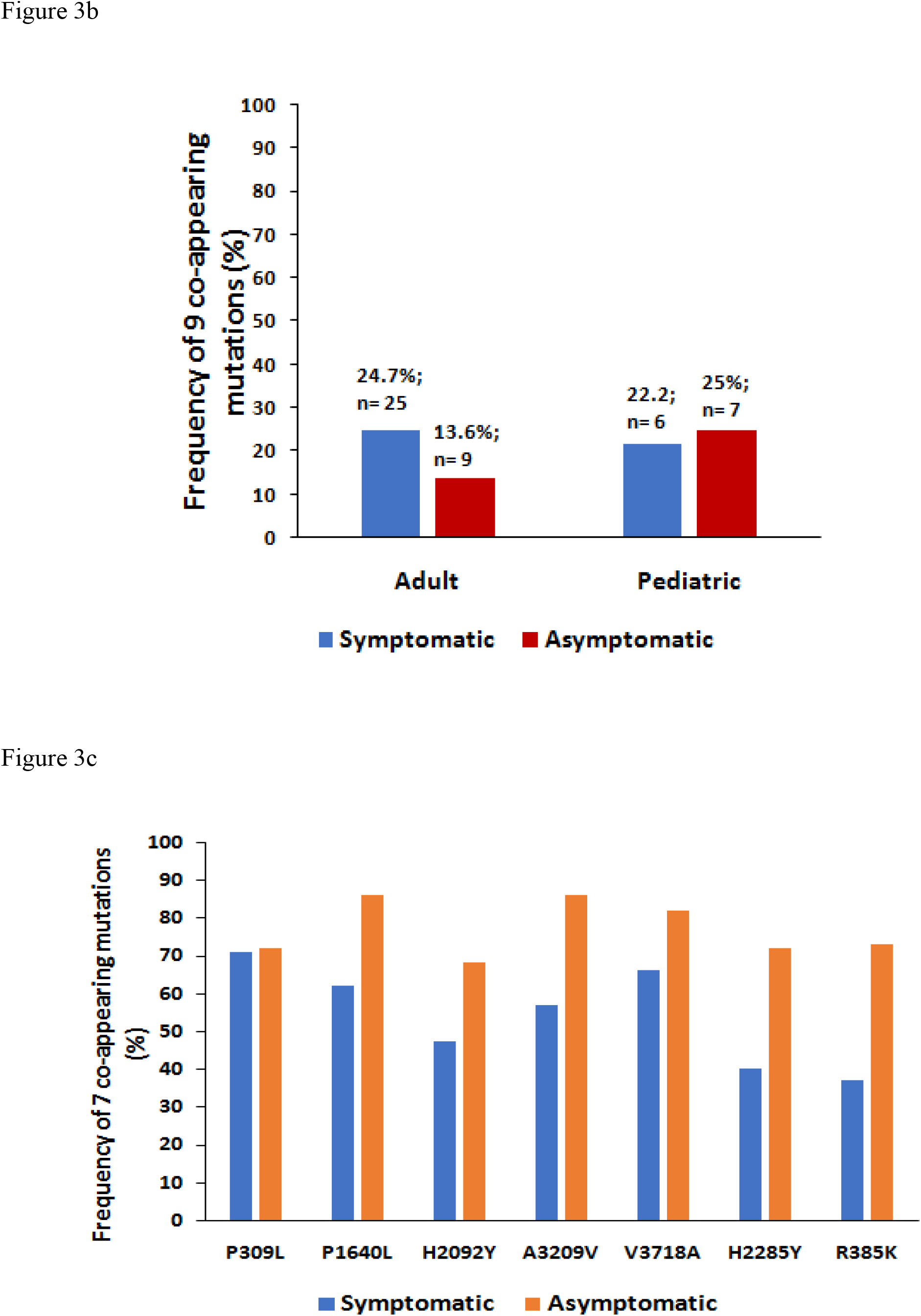
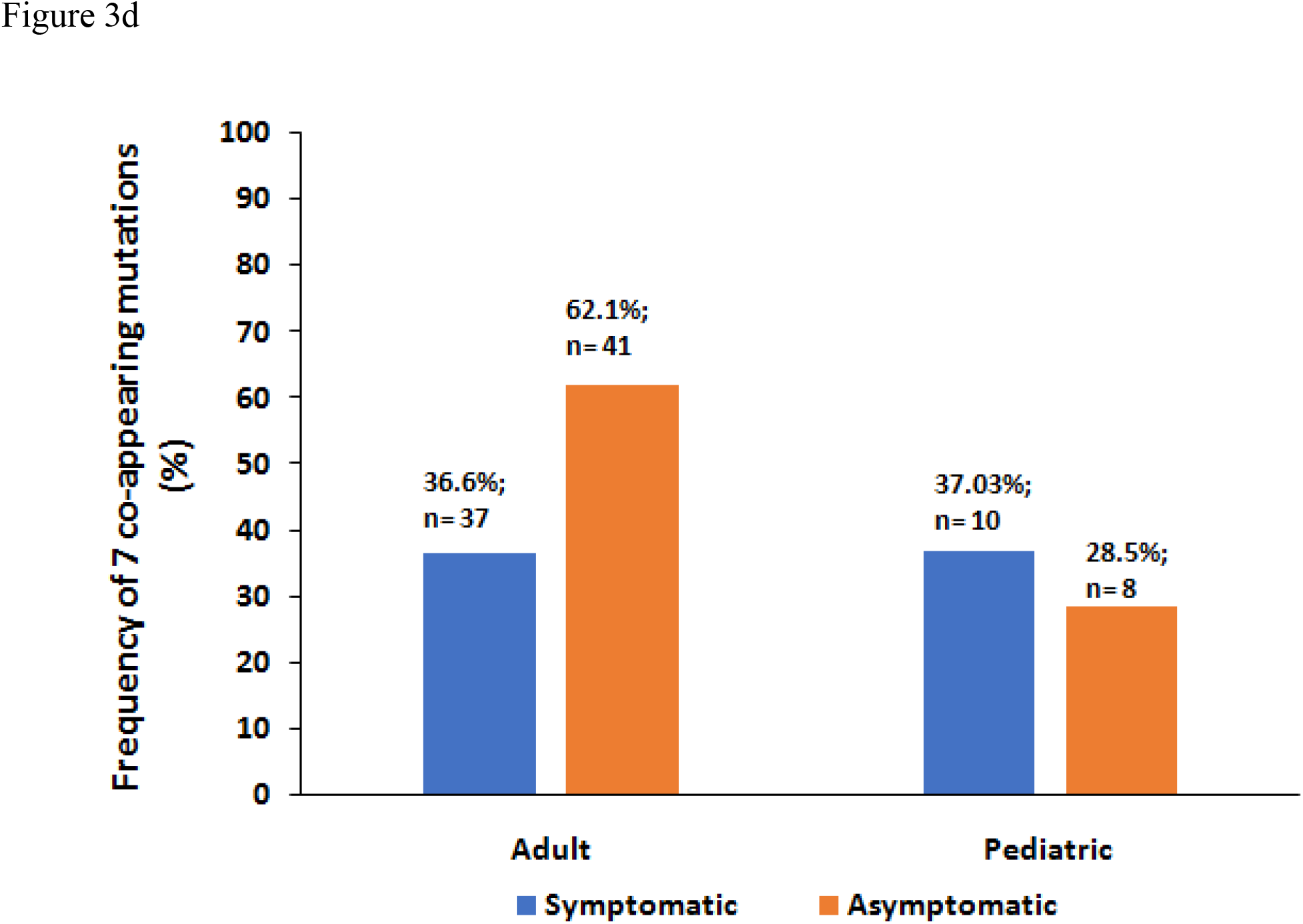
Frequency distribution of two mutually exclusive signature co-appearing mutation patterns across SARS-CoV-2 non-Spike proteins. (a) Frequency of mutations those were associated with asymptomatic disease outcome among (b) pediatric and adult age groups; (c) Mutations those were associated with symptomatic disease outcome among (d) pediatric and adult age groups.

Another set of 7 signature co-existing mutations (viz., P309L in NSP2, P1640L and H2092Y in NSP3, A3209V in NSP4, V3718A in NSP6, H2285Y in NSP15 and R385K in Nucleocapsid) was seen to predominate (1.7 times higher) among the asymptomatic patients (62.1%, 41/66) in comparison to the symptomatic ones (36.6%, 37/101) (Figure 3c). In contrast, among the paediatric group, marginal differences in frequency were evident among asymptomatic and symptomatic ones (28.5% and 37.03%, respectively) (Figure 3d). Among the 7 co-existing mutations, 3 were common to the 21I Delta sub-clade viz., NSP3:P1640L, NSP4:A3209V, and NSP6:V3718A. However, strains harboring this array of 7 co-existing mutations were only 2.73% more frequent among asymptomatic-vaccinated individuals (77.01%), in contrast to the asymptomatic-unvaccinated ones (74.2%).

It appears that the 2 discrete sets of non-Spike signature co-appearing mutations arrays were associated with the differential infection status of patients (symptomatic and asymptomatic, respectively). Statistical test also revealed that the association was significant (p-value was found to be 0.0074, which was ≤ 0.05). Symptomatic infection was seen to prevail due to the set of “9 signature non-spike co-appearing mutations”, even in the post-vaccination scenario.

### 3.4. Impact of predictive interactions of SARS-CoV-2 mutated proteins and host proteins involved in inflammatory responses

Some host-proteins which were reported to be physically associated with SARS-CoV-2 proteins [19] and involved in inflammatory responses were selected for our study. SARS-CoV-2 proteins harboring the identified co-occurring mutations (NSP2, NSP3, NSP4, NSP6, NSP14, NSP15, ORF7b and Nucleocapsid) were docked with the selected human proteins. We obtained different global energy values (binding energy for the interacting proteins) during docking the original and mutant SARS-CoV-2 proteins with some host proteins involved in inflammatory responses, among the symptomatic and asymptomatic groups. More negative value of binding energy represents more binding affinity between two proteins. 1.7 to 2-fold change of global energy among host protein(s) and the original or mutant viral proteins were considered to be significant in some previous studies [23]. While, in our analyses, we have highlighted all changes in global energy/binding energy of 1.5-fold or higher as significant.

Among the asymptomatic group, the mutant form of viral non-structural protein NSP2 (mutated site: P309L) revealed decreased binding affinity for host proteins like EIF4E2 (more global binding energy of mutant form, i.e., -28.34 than the original one, i.e., -42.90) and RAP1GDS1 (more global binding energy in mutant protein, i.e., -37.01 than the original form, i.e., -64.23) in comparison to the original protein. On the contrary, the affinity of mutant NSP2 was augmented in case of GIGYF2 protein (less global binding energy in mutant protein, i.e., -43.02 than the original form, i.e., -20.60). Regarding RNF41 host-protein which plays a significant role in suppressing the production of proinflammatory cytokines, though the fold increase is below 1.5 (i.e.1.39) it has been shown to interact with NSP15, and the binding affinity has been elevated in case of mutant NSP15 (H2285Y) compared to the original one. Also, for cellular proteins which are known to interact with SARS-CoV-2 Nucleocapsid, when mutations like R385K are present, the binding affinity is higher for cellular proteins like G3BP2, LARP1 (less global binding energy) and lower for G3BP1 (more global binding energy) (Table-2a).

**Table : 2a.**
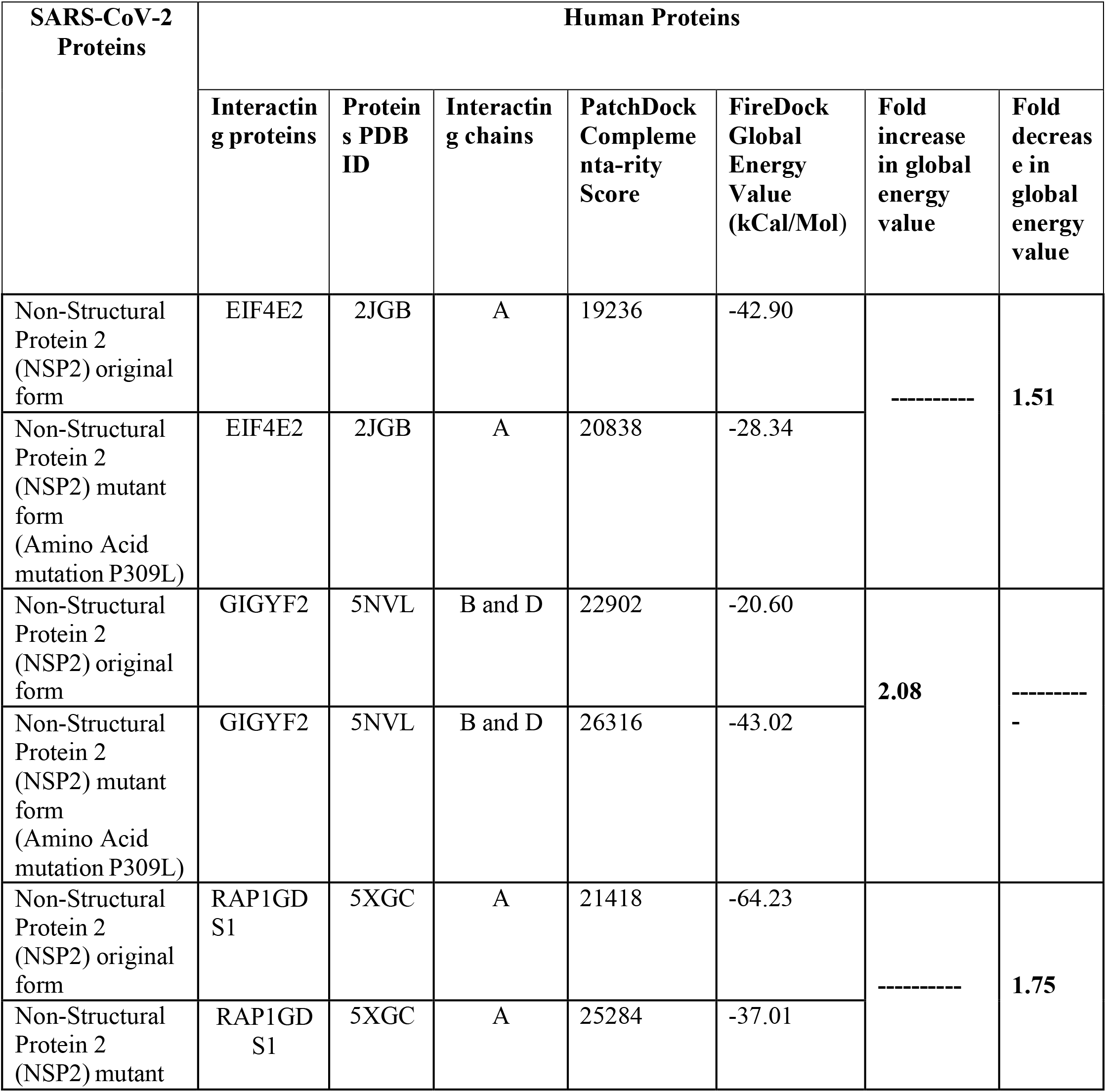
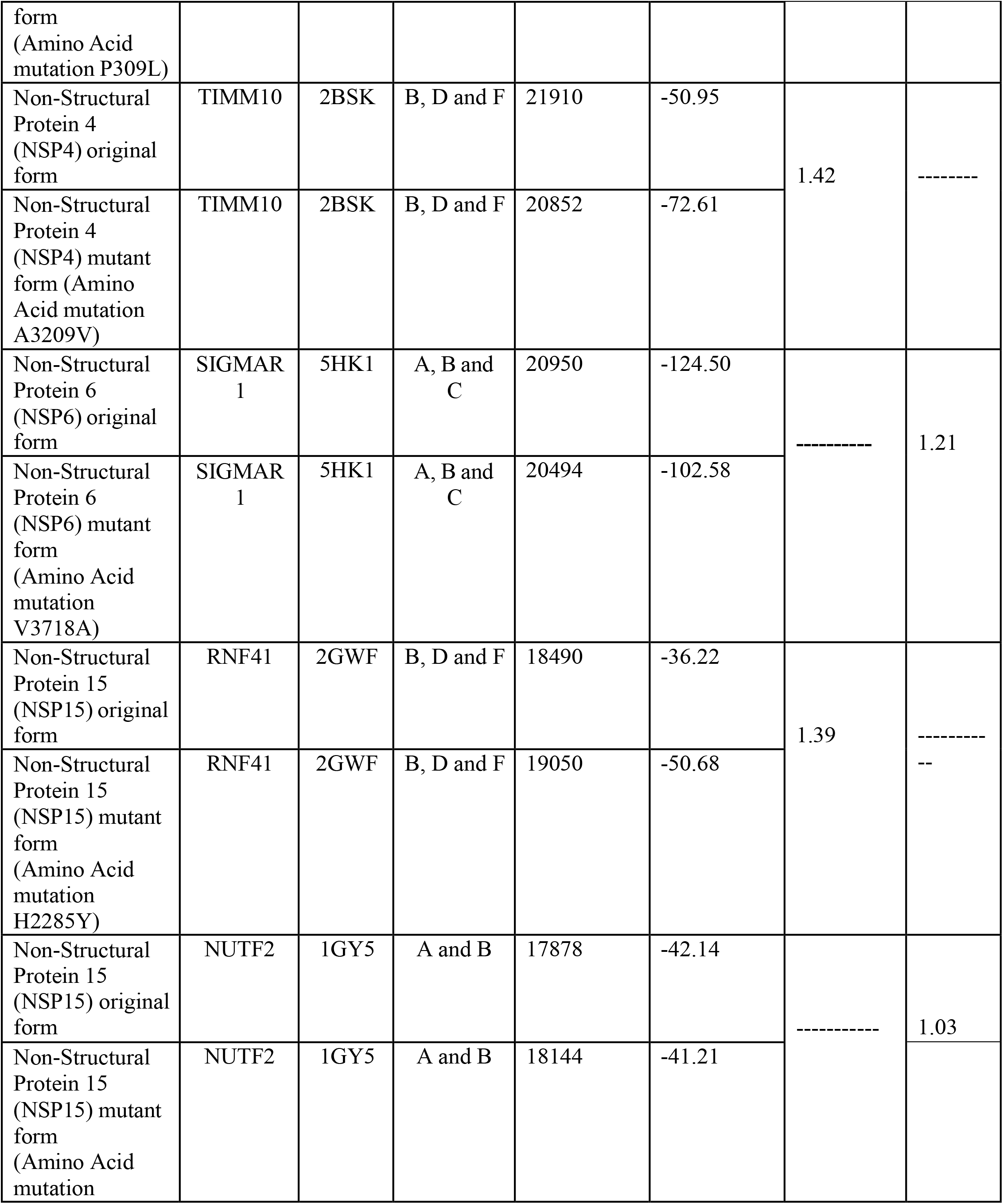
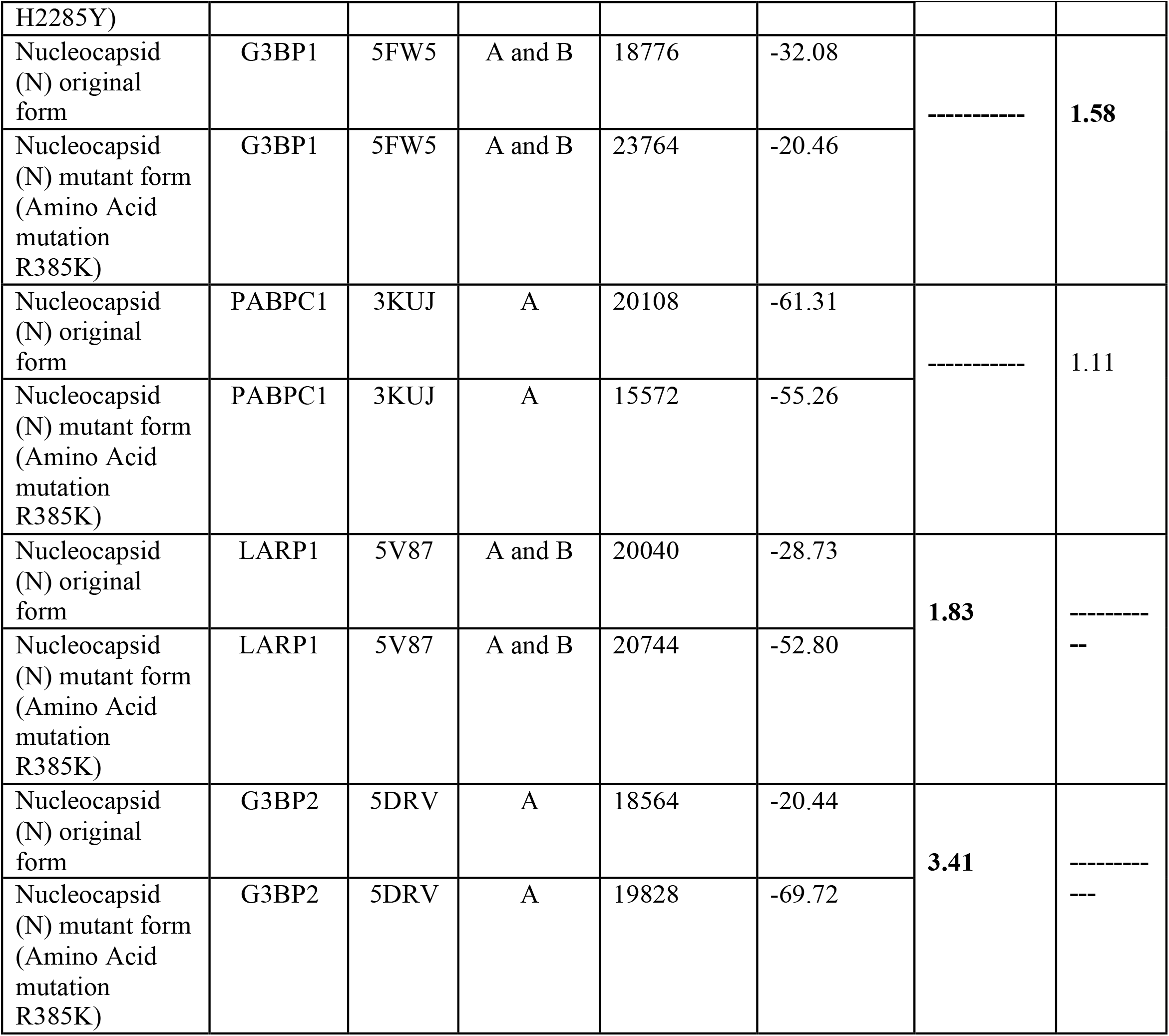
Analysis of fold change in global energy value regarding protein-protein interaction between host interacting proteins with the original and mutant SARS-CoV-2 proteins among asymptomatic patients. Values above 1.5 have been highlighted in **bold**. [Fold Increase = (global energy value of mutant SARS-CoV-2 protein/ global energy value of original SARS-CoV-2 protein), and Fold Decrease = 1/ (global energy value of mutant SARS-CoV-2 protein/ global energy value of original SARS-CoV-2 protein)]

Among all the co-occurring mutations in symptomatic patients, only Nucleocapsid protein revealed altered binding affinity to the specific host-proteins upon accumulation of mutations like G215C. Mutant N showed lower binding affinity to PABPC1 (more global binding energy in mutant protein, i.e., -36.63 than the original one, i.e., -61.31) (Table-2b). G215C mutation in N protein of symptomatic patients brought about 1.61-fold increase in binding affinity to G3BP2, which is much lower than what R385K mutation in N depicted in case of asymptomatic patients (3.41-fold increase in binding affinity).

**Table : 2b.**
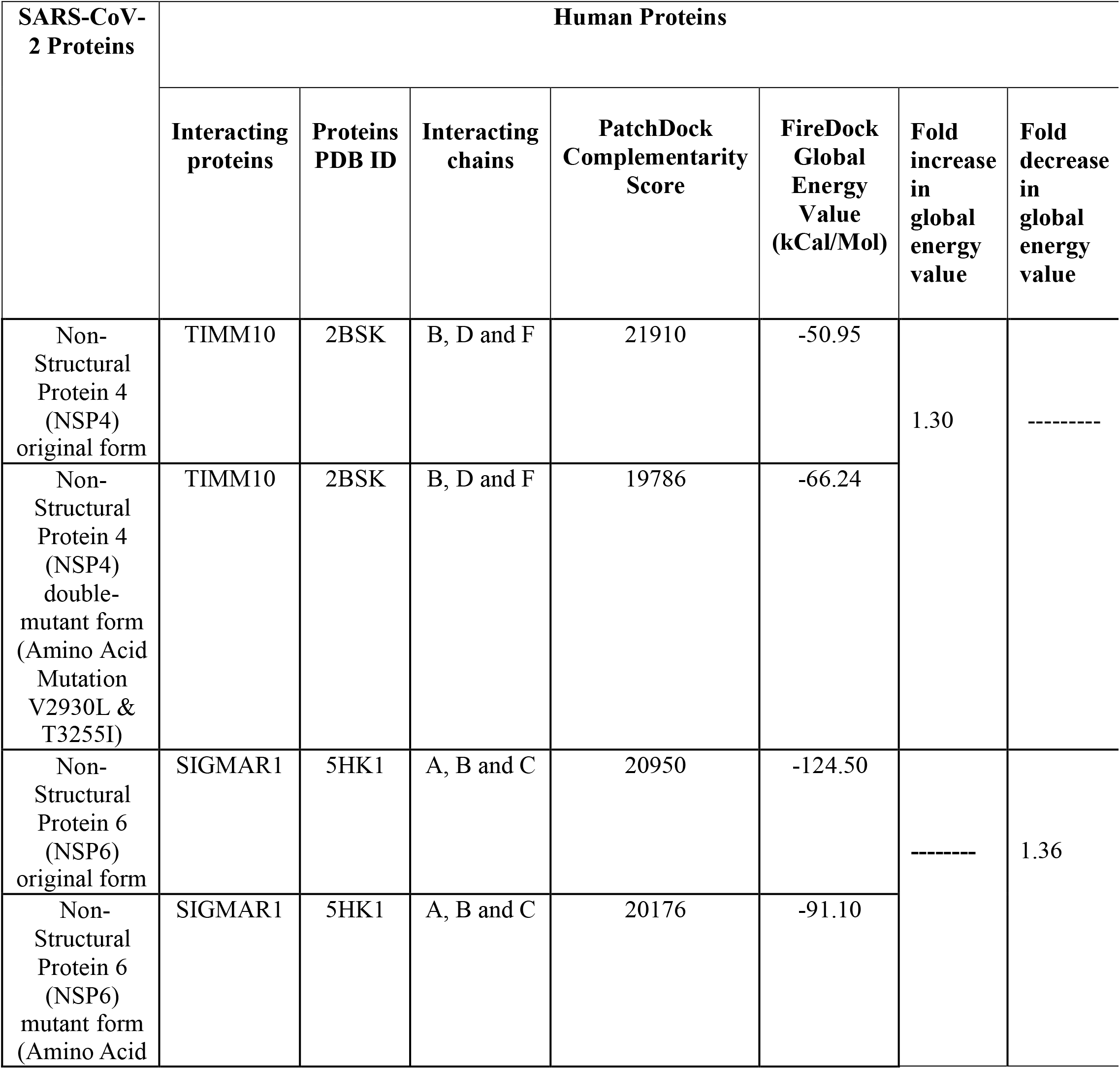
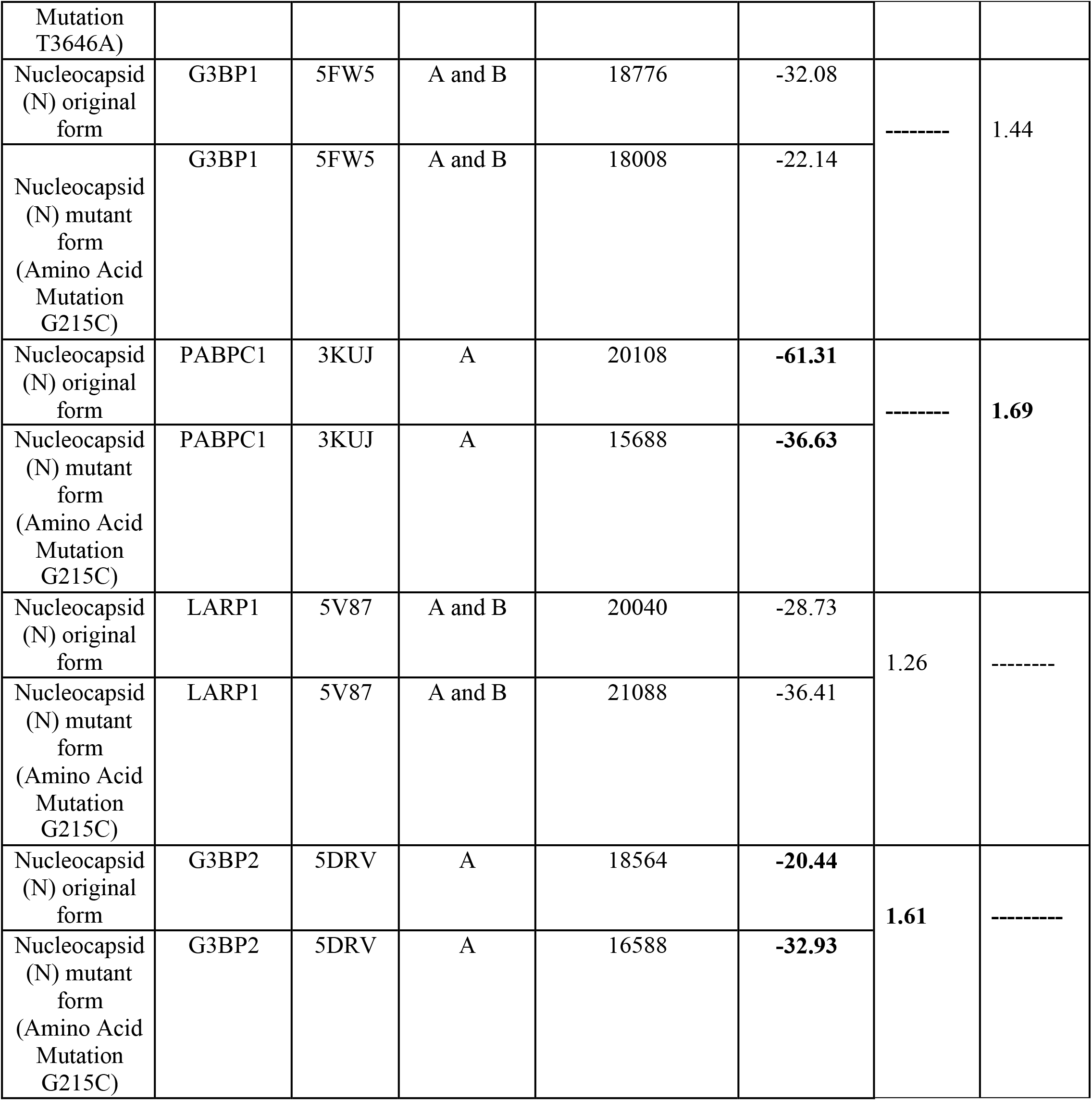
Analysis of fold change in global energy value regarding protein-protein interaction between host interacting proteins with original and mutant SARS-CoV-2 proteins among symptomatic patients. Values above 1.5 have been highlighted in **bold**. [Fold Increase = (global energy value of mutant SARS-CoV-2 protein/ global energy value of original SARS-CoV-2 protein), and Fold Decrease = 1/ (global energy value of mutant SARS-CoV-2 protein/ global energy value of original SARS-CoV-2 protein)]

Winding up, altered binding affinity of viral mutant proteins with host proteins engaged in inflammatory response could be one of the significant players behind symptomatic disease outcome.

## 4. Discussion

SARS-CoV-2 whole-genome sequencing provides an in-depth understanding of the mutation assemblage and phylodynamics of this deadly RNA virus [24-25]. The accumulation of mutations in RNA viruses like influenza virus revealed emergence of vaccine escape or drug-resistant mutants, demanding for a continuous need to design newer vaccines or therapeutics [26-27]. During the second COVID-19 wave in India, emergence of new mutations could have driven the higher transmissibility and pathogenicity of Delta VoC [28]. In addition to Spike mutations, the transpiring co-existing mutations within non-Spike proteins might also impact the efficacy of leading Spike-recombinant vaccines. Hence, tracking the continuously evolving SARS-CoV-2 mutations in samples in real-time can underscore their role in neutralization efficiency or immune evasion potential [29]. Nevertheless, our report is the first study from eastern India that portrayed a comparative scenario of co-appearing mutations encompassing the non-Spike proteins, and has associated them to the disease outcome among the vaccinated and unvaccinated patients.

Our study highlighted that the Delta VoC was the master player driving both symptomatic as well as asymptomatic infections during the second pandemic wave across eastern India, both in adult and pediatric patients, irrespective of the vaccination status. Delta was first detected in India in late 2020 and revealed itself to be extremely diffusive, worldwide (ECDC, 2021) [30]. Italso determined the sudden peaks of infections in multiple countries, even after the graph declined on account of massive vaccination drives [31-32]. Studies revealed that signature mutations across the Delta genome augmented its transmissibility and lowered the affinity for the neutralizing antibodies against previous variants [33-34]. We could identify 2 unique signature sets of co-existing mutations within Delta, which were mutually exclusive among the symptomatic and asymptomatic patients. Phylogeny revealed that the symptomatic strains were Delta sub-clade 21J-like, whereas the asymptomatic strains formed a completely new sub-cluster in between Delta 21A and Delta sub-clade 21I strains. These types of unique strains were also submitted to GISAID from a few other countries, but the GISAID metadata (submitted till 24^th^ January, 2022) revealed that they might have emerged in India prior to other countries.

Previous studies classified the Delta variant in two distinct groups: with and without the Nucleotide substitution G215C. Since the end of June 2021, the incidence of N: G215C Delta up-surged to about 95%, worldwide. In India, when vaccination was low during June 2021, the two strains of Delta reached similar levels, both occupying almost 50% of the total genomes [35]. Recently, few studies have highlighted the role of Nucleocapsid mutations in promoting higher infectivity and virulence of SARS-CoV-2 [36]. It was not only the Spike variations alone which allowed Delta to out-compete other variants in the second wave, rather the role of other non-Spike mutations have been underexplored to date [37-38]. N: G215C was one of the co-appearing mutations associated with post-vaccination symptomatic infection in our analyses. Significant variations across Nucleocapsid, ORF1a, ORF1b and ORF7b caused the divergence ofsub-clade 21J and 21I from the prototype 21A Delta. This might be a consequence of selection pressure due to growing vaccination rates.

We have attempted to extrapolate our unique observation within Delta VoC across this endemic setting to other emerging SARS-CoV-2 variants circulating worldwide (Figure 4). The T3255I mutation in NSP4 protein was selected in Lambda and Mu VoI and also in the recently emerged VoC (21K) Omicron [39-40]. Even the IHU variant (B.1.640.2) which has recently emerged in France harbors this T3255I mutation, generating symptomatic disease outcome [41]. Studies revealed that SARS-CoV-2 NSP4 protein interacts with different host-factors involved in antiviral innate immune signaling [42]. NSP4: T3255I mutation could have probably altered this interactome to evade the antiviral response, leading to better viral propagation and infectivity as evident in Omicron and in IHU [43]. Interestingly, the IHU variant also harbors NSP2: P309L and NSP6: V3718A mutations that were present in our asymptomatic signature co-occurring mutations array. Hence, IHU is the first SARS-CoV-2 variant where coexistence of both symptomatic (T3255I) and asymptomatic (P309L and V3718A) mutations has been evidenced. Therefore, it will be interesting to track its pathogenicity and diffusivity in contrast to other prevailing variants.

**Figure 4.**
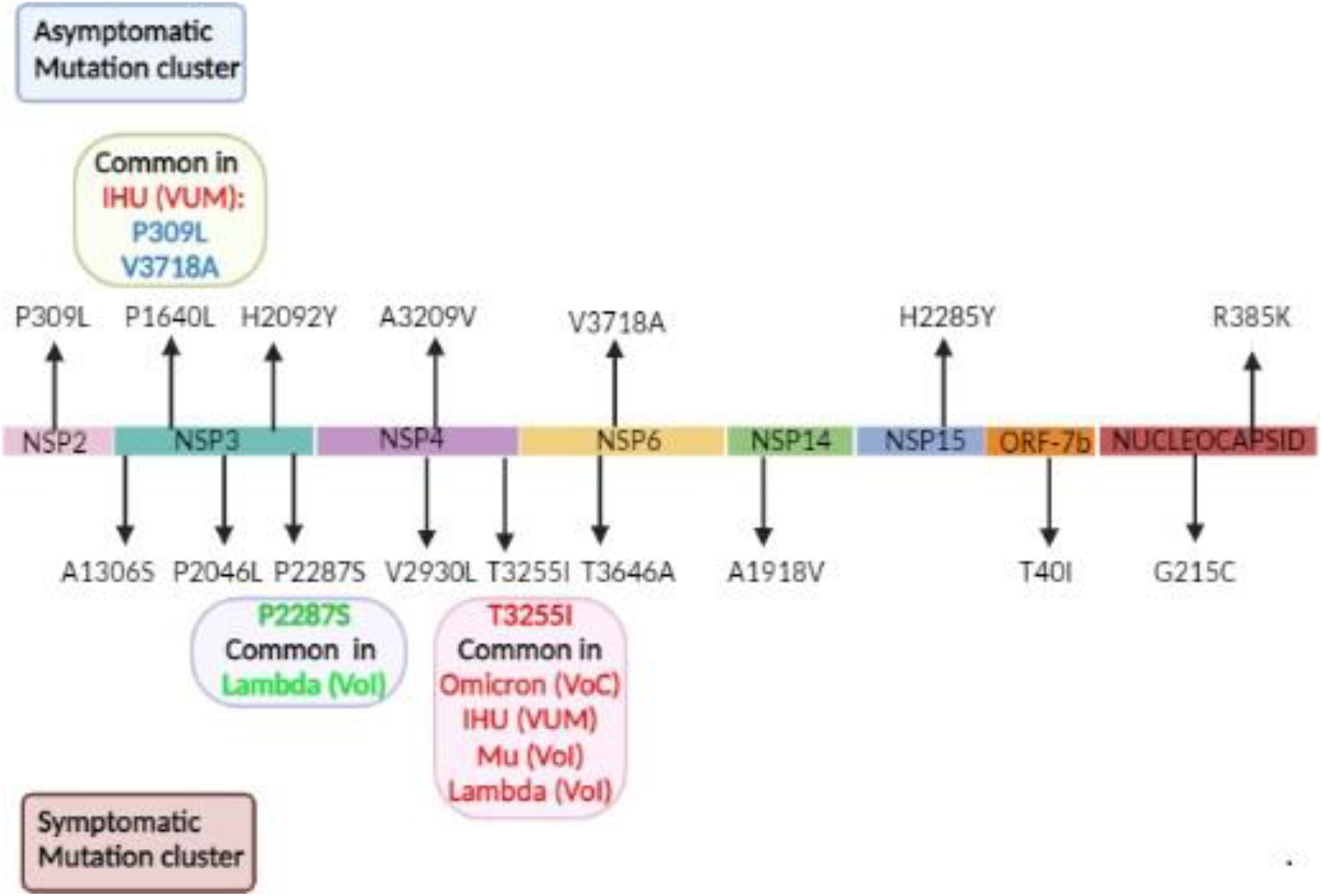
Schematic representation of signature co-appearing mutation clusters causing asymptomatic and symptomatic COVID-19 disease outcome. The prevalence and/or emergence of some of these mutations in other SARS-CoV-2 variants were also shown.

Among the set of 9 co-appearing mutations driving symptomatic infections, P2046L and P2287S within NSP3 were additional to the mutations present in 21J strains. Few reports suggest the de-mono-ADP-ribosylation (de-MARylation) of STAT1 by the SARS-CoV-2 NSP3 may be responsible for the cytokine storm observed in the most severe cases of COVID-19 [44]. The co-existence of P2046L and P2287S in NSP3 protein suggests their plausible role in augmenting the cytokine response which needs further experimental validation. This is indicative of the fact that adaptive pressure is leading to rapid SARS-CoV-2 evolution through emergence of recombinant clades within an endemic setting. It is highly probable that the strains harboring 9 signature co-appearing mutations array might give rise to a new virulent lineage in future. Asymptomatic infection is a significant factor driving the rapid spread of COVID-19 pandemic [45]. Due to the high risk of these silent spreaders, it is imperative to monitor the genomic constellation of the strains causing asymptomatic infections [46].

Mere association of specific viral mutations may not be an adequate predictor of disease severity. Host immuno-genetic determinants, comorbidities status and the treatment regime also decide the disease outcome. SARS-CoV-2 emerging variants imply multiple strategies to evade host immune responses by hijacking host proteins for their efficient replication [47]. The emergenceof highly transmissible and pathogenic 21J Delta with N: G215C mutation among populations with high vaccination rates should alert the health-care authorities regarding future management plans for the pandemic [48]. Recent studies reported that Nucleocapsid protein causes impairment of stress granule formation by targeting G3BP1 and G3BP2 proteins, which also playa role in antiviral signaling [49]. Through our analyses, we showed that G215C mutant N proteinimparted lower binding affinity to G3BP2 in symptomatic strains, in contrast to R385K mutant Nof asymptomatic strains. Further experimental validations are required to prove whether altered affinity of mutant N to G3BP2 could compromise its antiviral properties and give way to symptomatic infection.

## 5. Conclusion

We have identified two significant “John Hancock’’ sets of non-Spike co-appearing mutations within Delta and have traced their plausible emergence in India. This study is one of the first which has highlighted the correlation of these signature co-mutation constellations with differential COVID-19 infection status, even among the vaccinated patients. The selection of some of these mutations in highly diffusive variants like Omicron and IHU underscores the importance of continuous surveillance of these signature mutation sets. The impact of these constellations of co-existing mutations on evasion of current vaccine-induced immunity requires further in-depth experimental analysis and clinical evidence.

## Acknowledgements

The authors acknowledge financial and overall support provided by the Department of Biotechnology, Ministry of Science and Technology, India, and Department of Health Research/ Indian Council of Medical Research, India. S Das would like to acknowledge support from J C Bose fellowship. AB would like to acknowledge support from Science and Engineering Research Board-Department of Science and Technology. JR would like to acknowledge support from University Grants Commission.

The scientific and technical staff working at Regional VRDL at ICMR-National Institute of Cholera and Enteric Diseases (Ashis Debnath, Pradip Kumar Jana, Soumen Mukherjee, Rudrak Gupta, Madhumonti Biswas, Sutapa Hazra, Chinmoy Mondal, Satyabrata Ghorai, Souvik Kar, Asish Kumar Jana, Musarraf Hossain, Kartick Chandra Mondal, Avisek Sinha, Biswajit Dey, Nayan Basuli) are acknowledged for their contribution in processing of nasopharyngeal and oropharyngeal swab samples and determination of SARS-CoV-2 positivity. The National Institute of BioMedical Genomics COVID-19 Sequencing laboratory members (Shreelekha Dutta, Subrata Patra, Trinath Ghosh, Sumanta Sarkar, Shekhar Ghosh, Debojyoti Roy, Sabyasachi Bhattacharya, Meghna Chowdhury, Surajit Mahapatra, Animesh Kumar Singh, Antara Paul, Rezwanuzzaman Laskar) are hereby acknowledged for their contribution in conducting the viral genome sequencing of the COVID-19 positive samples.

---

**Supplementary figure 1:**
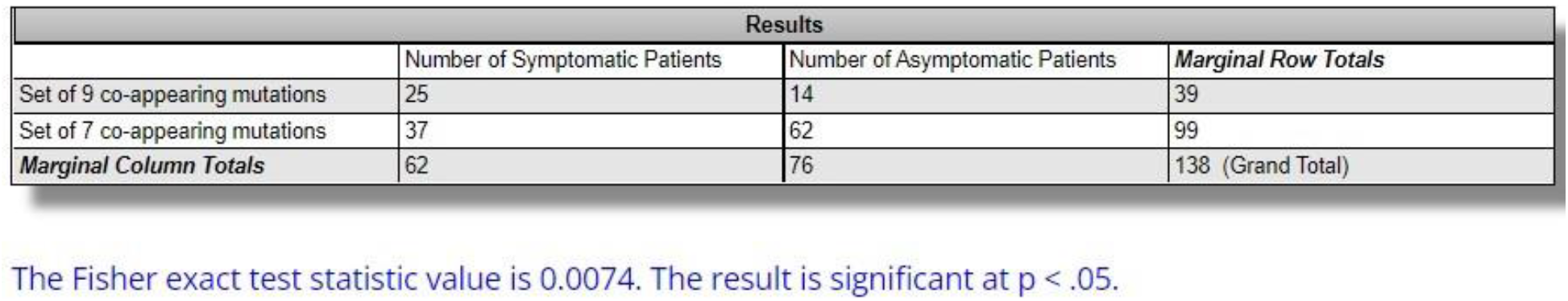
Fisher’s exact statistical test of the association of the 9 and 7 co-appearing mutations with respective clinical groups. Fisher exact statistical value of 0.0074 depicts that the result is significant at p value less than 0.05.

## References

1. Schiller, H.B., van Breugel, M. and Nawijn, M.C., 2021. SARS-CoV-2-specific hotspots in virus–host interaction networks. Nature immunology, pp.1–3. https://doi.org/10.1038/s41590-021-00963-9

2. Wang, R., Chen, J. and Wei, G.W., 2021. Mechanisms of sars-cov-2 evolution revealing vaccine-resistant mutations in europe and america. The journal of physical chemistry letters, 12, pp.11850–11857. https://doi.org/10.1021/acs.jpclett.1c03380

3. Harvey, W.T., Carabelli, A.M., Jackson, B., Gupta, R.K., Thomson, E.C., Harrison, E.M., Ludden, C., Reeve, R., Rambaut, A., Peacock, S.J. and Robertson, D.L., 2021. SARS-CoV-2 variants, spike mutations and immune escape. Nature Reviews Microbiology, 19(7), pp.409–424. https://doi.org/10.1038/s41579-021-00573-0

4. Planas, D., Veyer, D., Baidaliuk, A., Staropoli, I., Guivel-Benhassine, F., Rajah, M.M., Planchais, C., Porrot, F., Robillard, N., Puech, J. and Prot, M., 2021. Reduced sensitivity of SARS-CoV-2 variant Delta to antibody neutralization. Nature, 596(7871), pp.276–280. https://doi.org/10.1038/s41586-021-03777-9

5. Gandhi, R.T., Lynch, J.B. and Del Rio, C., 2020. Mild or moderate Covid-19. New England Journal of Medicine, 383(18), pp.1757–1766. https://doi.org/10.1038/s41586-021-03944-y

6. Gandhi, R.T., Lynch, J.B. and Del Rio, C., 2020. Mild or moderate Covid-19. New England Journal of Medicine, 383(18), pp.1757–1766. https://doi:10.1056/NEJMcp2009249

7. Brodin, P., 2021. Immune determinants of COVID-19 disease presentation and severity. Nature Medicine, 27(1), pp.28–33. https://doi.org/10.1038/s41591-020-01202-8

8. Asrani, P., Eapen, M.S., Hassan, M.I. and Sohal, S.S., 2021. Implications of the second wave of COVID-19 in India. The Lancet Respiratory Medicine, 9(9), pp.e93–e94. https://doi.org/10.1016/S2213-2600(21)00312-X

9. Rowley, A.H., 2020. Understanding SARS-CoV-2-related multisystem inflammatory syndrome in children. Nature Reviews Immunology, 20(8), pp.453–454. https://doi.org/10.1038/s41577-020-0367-5

10. Biswas, N., Mallick, P., Maity, S.K., Bhowmik, D., Mitra, A.G., Saha, S., Roy, A., Chakrabarti, P., Paul, S. and Chakrabarti, S., 2021. Genomic surveillance and phylodynamic analyses reveal emergence of novel mutation and co-mutation patterns within SARS-CoV2 variants prevalent in India. bioRxiv. https://doi.org/10.3389/fmicb.2021.703933

11. Barton, M.I., MacGowan, S.A., Kutuzov, M.A., Dushek, O., Barton, G.J. and van der Merwe, P.A., 2021. Effects of common mutations in the SARS-CoV-2 Spike RBD and its ligand, the human ACE2 receptor on binding affinity and kinetics. Elife, 10, p.e70658. https://doi.org/10.7554/elife.70658

12. Greaney, A.J., Starr, T.N., Gilchuk, P., Zost, S.J., Binshtein, E., Loes, A.N., Hilton, S.K., Huddleston, J., Eguia, R., Crawford, K.H. and Dingens, A.S., 2021. Complete mapping of mutations to the SARS-CoV-2 spike receptor-binding domain that escape antibody recognition. Cell host & microbe, 29(1), pp.44–57. https://doi.org/10.1016/j.chom.2020.11.007

13. Wu, H., Xing, N., Meng, K., Fu, B., Xue, W., Dong, P., Tang, W., Xiao, Y., Liu, G., Luo, H. and Zhu, W., 2021. Nucleocapsid mutations R203K/G204R increase the infectivity, fitness, and virulence of SARS-CoV-2. Cell host & microbe, 29(12), pp.1788–1801. https://doi.org/10.1016/j.chom.2021.11.005

14. Wu, S., Tian, C., Liu, P., Guo, D., Zheng, W., Huang, X., Zhang, Y. and Liu, L., 2021. Effects of SARS-CoV-2 mutations on protein structures and intraviral protein–protein interactions. Journal of medical virology, 93(4), pp.2132–2140. https://doi.org/10.1002/jmv.26597

15. McCoy, K., Peterson, A., Tian, Y. and Sang, Y., 2020. Immunogenetic association underlying severe COVID-19. Vaccines, 8(4), p.700. https://doi.org/10.3390/vaccines8040700

16. Baggen, J., Vanstreels, E., Jansen, S. and Daelemans, D., 2021. Cellular host factors for SARS-CoV-2 infection. Nature Microbiology, 6(10), pp.1219–1232. https://doi.org/10.1038/s41564-021-00958-0

17. Karaderi, T., Bareke, H., Kunter, I., Seytanoglu, A., Cagnan, I., Balci, D., Barin, B., Hocaoglu, M.B., Rahmioglu, N., Asilmaz, E. and Taneri, B., 2020. Host genetics at the intersection of autoimmunity and COVID-19: A potential key for heterogeneous COVID-19 severity. Frontiers in Immunology, 11, p.3314. https://doi.org/10.3389/fimmu.2020.586111

18. Sungnak, W., Huang, N., Bécavin, C., Berg, M., Queen, R., Litvinukova, M., Talavera-López, C., Maatz, H., Reichart, D., Sampaziotis, F. and Worlock, K.B., 2020. SARS-CoV-2 entry factors are highly expressed in nasal epithelial cells together with innate immune genes. Nature medicine, 26(5), pp.681–687. https://doi.org/10.1038/s41591-020-0868-6

19. Gordon, D.E., Jang, G.M., Bouhaddou, M., Xu, J., Obernier, K., White, K.M., O’Meara, M.J., Rezelj, V.V., Guo, J.Z., Swaney, D.L. and Tummino, T.A., 2020. A SARS-CoV-2 protein interaction map reveals targets for drug repurposing. Nature, 583(7816), pp.459–468. https://doi.org/10.1038/s41586-020-2332-7

20. Wang, S., Li, W., Liu, S. and Xu, J., 2016. RaptorX-Property: a web server for protein structure property prediction. Nucleic acids research, 44(W1), pp.W430–W435. https://doi.org/10.1093/nar/gkw306

21. Schneidman-Duhovny, D., Inbar, Y., Nussinov, R. and Wolfson, H.J., 2005. PatchDock and SymmDock: servers for rigid and symmetric docking. Nucleic acids research, 33(suppl_2), pp.W363–W367. https://doi.org/10.1093/nar/gki481

22. Fornes, O., Garcia-Garcia, J., Bonet, J. and Oliva, B., 2014. On the use of knowledge-based potentials for the evaluation of models of protein–protein, protein–DNA, and protein–RNA interactions. Advances in protein chemistry and structural biology, 94, pp.77–120. http://dx.doi.org/10.1093/nar/gki481

23. Kumar, A. and Ramanathan, K., 2015. Virtual screening approach to identify potential ALK inhibitor from traditional Chinese medicine database. Research Journal of Pharmaceutical, Biological and Chemical Sciences, 6(1), pp.94–101.

24. Maitra, A., Sarkar, M.C., Raheja, H., Biswas, N.K., Chakraborti, S., Singh, A.K., Ghosh, S., Sarkar, S., Patra, S., Mondal, R.K. and Ghosh, T., 2020. Mutations in SARS-CoV-2 viral RNA identified in Eastern India: Possible implications for the ongoing outbreak in India and impact on viral structure and host susceptibility. Journal of Biosciences, 45(1), pp.1–18. https://doi.org/10.1007/s12038-020-00046-1

25. Zhang, Y.Z. and Holmes, E.C., 2020. A genomic perspective on the origin and emergence of SARS-CoV-2. Cell, 181(2), pp.223–227. https://doi.org/10.1016/j.cell.2020.03.035

26. Zharikova, D., Mozdzanowska, K., Feng, J., Zhang, M. and Gerhard, W., 2005. Influenza type A virus escape mutants emerge in vivo in the presence of antibodies to the ectodomain of matrix protein 2. Journal of virology, 79(11), pp.6644–6654. https://doi.org/10.1128/jvi.79.11.6644-6654.2005

27. Smirnov, Y.A., Gitelman, A.K., Govorkova, E.A., Lipatov, A.S. and Kaverin, N.V., 2004. Influenza H5 virus escape mutants: immune protection and antibody production in mice. Virus research, 99(2), pp.205–208. https://doi.org/10.1016/j.virusres.2003.11.012

28. Pouwels, K.B., Pritchard, E., Matthews, P.C., Stoesser, N., Eyre, D.W., Vihta, K.D., House, T., Hay, J., Bell, J.I., Newton, J.N. and Farrar, J., 2021. Effect of Delta variant on viral burden and vaccine effectiveness against new SARS-CoV-2 infections in the UK. Nature medicine, 27(12), pp.2127–2135. https://doi.org/10.1038/s41591-021-01548-7

29. Grant, R., Charmet, T., Schaeffer, L., Galmiche, S., Madec, Y., Von Platen, C., Chény, O., Omar, F., David, C., Rogoff, A. and Paireau, J., 2021. Impact of SARS-CoV-2 Delta variant on incubation, transmission settings and vaccine effectiveness: Results from a nationwide case-control study in France. The Lancet Regional Health-Europe, p.100278. https://doi.org/10.1016/j.lanepe.2021.100278

30. Kannan, S.R., Spratt, A.N., Cohen, A.R., Naqvi, S.H., Chand, H.S., Quinn, T.P., Lorson, C.L., Byrareddy, S.N. and Singh, K., 2021. Evolutionary analysis of the Delta and Delta Plus variants of the SARS-CoV-2 viruses. Journal of autoimmunity, 124, p.102715. https://doi.org/10.1016/j.jaut.2021.102715

31. Mattiuzzi, C., Henry, B.M. and Lippi, G., 2021. Is diffusion of SARS-CoV-2 variants of concern associated with different symptoms?: Symptoms of COVID-19. The Journal of Infection. https://doi:10.1016/j.jinf.2021.07.008

32. Shastri, J., Parikh, S., Aggarwal, V., Agrawal, S., Chatterjee, N., Shah, R., Devi, P., Mehta, P. and Pandey, R., 2021. Severe SARS-CoV-2 breakthrough reinfection with delta variant after recovery from breakthrough infection by alpha variant in a fully vaccinated health worker. Frontiers in Medicine, p.1379. https://doi.org/10.3389/fmed.2021.737007

33. McCallum, M., Walls, A.C., Sprouse, K.R., Bowen, J.E., Rosen, L.E., Dang, H.V., De Marco, A., Franko, N., Tilles, S.W., Logue, J. and Miranda, M.C., 2021. Molecular basis of immune evasion by the Delta and Kappa SARS-CoV-2 variants. Science, p.eabl8506. https://doi.org/10.1126/science.abl8506

34. Kuzmina, A., Wattad, S., Khalaila, Y., Ottolenghi, A., Rosental, B., Engel, S., Rosenberg, E. and Taube, R., 2021. SARS CoV-2 Delta variant exhibits enhanced infectivity and a minor decrease in neutralization sensitivity to convalescent or post-vaccination sera. Iscience, 24(12), p.103467. https://doi.org/10.1016/j.isci.2021.103467

35. Marchitelli, V., Troise, C., Parisi, A., Bianco, A., Del Sambro, L., Somma, R., Coviello, A., Harabaglia, P., Ungaro, F. and De Natale, G., 2021. Evidence for the dependence of the SARS-Cov-2 Delta high diffusivity on the associated N: G215C nucleocapsid mutation. https://doi.org/10.21203/rs.3.rs-846225/v1

36. Wu, H., Xing, N., Meng, K., Fu, B., Xue, W., Dong, P., Tang, W., Xiao, Y., Liu, G., Luo, H. and Zhu, W., 2021. Nucleocapsid mutations R203K/G204R increase the infectivity, fitness, and virulence of SARS-CoV-2. Cell host & microbe, 29(12), pp.1788–1801. https://doi.org/10.1101/2021.05.24.445386

37. Burbelo, P.D., Riedo, F.X., Morishima, C., Rawlings, S., Smith, D., Das, S., Strich, J.R., Chertow, D.S., Davey Jr, R.T. and Cohen, J.I., 2020. Sensitivity in detection of antibodies to nucleocapsid and spike proteins of severe acute respiratory syndrome coronavirus 2 in patients with coronavirus disease 2019. The Journal of infectious diseases, 222(2), pp.206–213. https://doi.org/10.1093/infdis/jiaa273

38. Brochot, E., Demey, B., Touzé, A., Belouzard, S., Dubuisson, J., Schmit, J.L., Duverlie, G., Francois, C., Castelain, S. and Helle, F., 2020. Anti-spike, anti-nucleocapsid and neutralizing antibodies in SARS-CoV-2 inpatients and asymptomatic individuals. Frontiers in microbiology, 11, p.2468. https://doi.org/10.3389/fmicb.2020.584251

39. Kandeel, M., Mohamed, M.E.M., Abd El-Lateef, H.M., Venugopala, K.N. and El-Beltagi, H.S., 2021. Omicron variant genome evolution and phylogenetics. Journal of medical virology. https://doi.org/10.1002/jmv.27515

40. He, X., Hong, W., Pan, X., Lu, G. and Wei, X., 2021. SARS-CoV-2 Omicron variant: characteristics and prevention. MedComm. https://doi.org/10.1002/mco2.110

41. Colson, P., Delerce, J., Burel, E., Dahan, J., Jouffret, A., Fenollar, F., Yahi, N., Fantini, J., La Scola, B. and Raoult, D., 2021. Emergence in Southern France of a new SARS-CoV-2 variant of probably Cameroonian origin harbouring both substitutions N501Y and E484K in the spike protein. medRxiv. https://doi.org/10.1101/2021.12.24.21268174

42. Davies, J.P., Almasy, K.M., McDonald, E.F. and Plate, L., 2020. Comparative multiplexed interactomics of SARS-CoV-2 and homologous coronavirus nonstructural proteins identifies unique and shared host-cell dependencies. ACS infectious diseases, 6(12), pp.3174–3189. https://dx.doi.org/10.1021/acsinfecdis.0c00500?ref=pdf

43. Lupala, C.S., Ye, Y., Chen, H., Su, X.D. and Liu, H., 2021. Mutations on RBD of SARS-CoV-2 Omicron variant result in stronger binding to human ACE2 receptor. Biochemical and Biophysical Research Communications. https://doi.org/10.1016/j.bbrc.2021.12.079

44. Claverie, J.M., 2020. A putative role of de-mono-ADP-ribosylation of STAT1 by the SARS-CoV-2 Nsp3 protein in the cytokine storm syndrome of COVID-19. Viruses, 12(6), p.646. https://doi.org/10.3390/v12060646

45. Oran, D.P. and Topol, E.J., 2020. Prevalence of asymptomatic SARS-CoV-2 infection: a narrative review. Annals of internal medicine, 173(5), pp.362–367. https://doi.org/10.7326/M20-3012

46. Nagy, Á., Pongor, S. and Győrffy, B., 2021. Different mutations in SARS-CoV-2 associate with severe and mild outcome. International journal of antimicrobial agents, 57(2), p.106272. https://doi.org/10.1016/j.ijantimicag.2020.106272

47. McCallum, M., Walls, A.C., Sprouse, K.R., Bowen, J.E., Rosen, L.E., Dang, H.V., De Marco, A., Franko, N., Tilles, S.W., Logue, J. and Miranda, M.C., 2021. Molecular basis of immune evasion by the Delta and Kappa SARS-CoV-2 variants. Science, p.eabl8506. https://doi.org/10.1126/science.abl8506

48. Bager, P., Wohlfahrt, J., Rasmussen, M., Albertsen, M. and Krause, T., 2021. Hospitalisation associated with SARS-CoV-2 delta variant in Denmark. The Lancet. Infectious Diseases. https://doi.org/10.1016/S1473-3099(21)00580-6

49. Zheng, Z.Q., Wang, S.Y., Xu, Z.S., Fu, Y.Z. and Wang, Y.Y., 2021. SARS-CoV-2 nucleocapsid protein impairs stress granule formation to promote viral replication. Cell Discovery, 7(1), pp.1–11. https://doi.org/10.1038/s41421-021-00275-0

